# Deep functional synthesis: a machine learning approach to gene functional enrichment

**DOI:** 10.1101/824086

**Authors:** Sheng Wang, Jianzhu Ma, Samson Fong, Stefano Rensi, Jiawei Han, Jian Peng, Dexter Pratt, Russ B. Altman, Trey Ideker

## Abstract

Gene functional enrichment is a mainstay of genomics, but it relies on manually curated databases of gene functions that are incomplete and unaware of the biological context. Here we present an alternative machine learning approach, Deep Functional Synthesis (DeepSyn), which moves beyond gene function databases to dynamically infer the functions of a gene set from its associated network of literature and data, conditioned on the disease and drug context of the current experiment. Using a knowledge graph with 3,048,803 associations between genes, diseases, drugs, and functions, DeepSyn obtained accurate performance (range 0.74 AUC to 0.96 AUC) on a variety of biological applications including drug target identification, gene set functional enrichment, and disease gene prediction.

**Availability:** The DeepSyn codebase is available on GitHub at http://github.com/wangshenguiuc/DeepSyn/ under an open source distribution license.

## MAIN TEXT

A common outcome of genomic analysis is the discovery of gene sets underlying specific biological functions. For instance, gene expression analyses produce sets of genes that are differentially expressed across conditions, or that cluster by expression similarity. Proteomics experiments produce lists of proteins and, by implication, their encoding genes, and so on. In all of these cases, the basic hypothesis is that the identified genes work coherently towards the same biological processes or functions. To label these functions, one turns to functional enrichment analysis. Among the numerous approaches that have been developed (Rhee et al., 2008; Zhou et al., 2017), some of the more widely used ones are the hypergeometric statistic (Breitling et al., 2004; Huang et al., 2009; Pomaznoy et al., 2018; Zeeberg et al., 2003) and gene set enrichment analysis (GSEA) (Al-Shahrour et al., 2007; Backes et al., 2007; Beissbarth and Speed, 2004; Subramanian et al., 2005), which seek to identify overlaps between the identified set of genes and those from a separate, pre-defined catalog of gene sets associated with known biological functions and pathways (Cerami et al., 2011; Fabregat et al., 2018; Kanehisa and Goto, 2000; Pico et al., 2008; Wang et al., 2018a).

Paradoxically, a gene set for which there is a very strong functional enrichment may be of less interest to researchers, since the set and its function have already been well characterized by previous studies. Of greater interest are gene sets that fail functional enrichment, or overlap known functions only marginally, because it is precisely from these ‘failures’ that new biological findings emerge. In these cases, an immediate next step is to explore the biological literature, as well as complementary data sets, to learn as much as possible about the genes in question. The goal is to mine knowledge pertinent to each gene and then to use this knowledge to synthesize mechanistic hypotheses for a function that might be held in common by all genes in the set. As a result, new functional categories might be defined and added to existing collections of functions. This process of discerning relevant findings from data and literature, and reasoning on this information to synthesize functional hypotheses, has not been widely automated but is one of the central tasks performed by a genome scientist.

When reasoning about gene functions, it is critical to include knowledge of the relevant experimental conditions and biological contexts under which the gene set has been identified. For instance, FoxA family transcription factors have been found to display different roles in a very strong tissue-specific manner. FoxA regulates glucagon expression in the pancreas, GLUT2 expression in the liver, and tyrosine hydroxylase expression in dopaminergic neurons (Fox et al., 2013). Such knowledge is strikingly absent from gene set functional enrichment tools, since their computational models and databases cannot easily encode the practically infinite space of biological conditions.

Here, we sought to develop a new reasoning tool, Deep Functional Synthesis (DeepSyn), to learn the biological functions of a gene set from its totality of literature and data. Our goal was to achieve two key advances over functional enrichment: the ability to synthesize new functional hypotheses rather than rely on the predefined functions of predefined gene sets, and the ability to guide these functional hypotheses by relevant biological conditions.

Biomedical relation extraction (BRE) (Hristovski et al., 2003; Lever et al., 2019; Tsuruoka et al., 2011) can be viewed as a first attempt towards our goal. BRE is able to capture statistical associations between entities referenced in literature, such as a gene name and a disease name that are co-mentioned in many abstracts. DeepSyn extends this concept by integrating associations from literature with those from primary data, and by moving beyond individual associations to build a global knowledge graph in which to trace highly relevant pathways of associations among genes, functions, diseases and/or drugs (e.g. linking disruption of a gene to changes in the activity of proteins, pathways, and incidence of disease in relevant experimental datasets).

### Defining and embedding biological functions in a global knowledge graph

The DeepSyn knowledge graph currently captures 3,048,803 associations among 85,180 biological entities including 27,175 genes, 26,365 diseases, 4,125 drugs, and 27,515 functions (Figure 1A, **KEY RESOURCE TABLE**, **STAR Methods**). Gene, disease, and drug identities are defined from public databases according to standard nomenclature (Genes: HNSC; Diseases: MeSH; Drugs: Chemical name). Functions, which have been less standardized, are defined by a combination of public databases (8,542 Gene Ontology term names) and entity recognition from biomedical abstracts using a deep neural network model (yielding another 18,973 distinct biological functions). Associations among these entities are mined from biomedical abstracts (e.g. probability of co-occurrence among genes and functions or pairs of functions), ontologies (e.g. associating drugs with targets, genes with diseases) or experimental databases (e.g. associating genes via protein-protein interactions or genes and diseases through genome-wide association studies).

**Figure 1.**
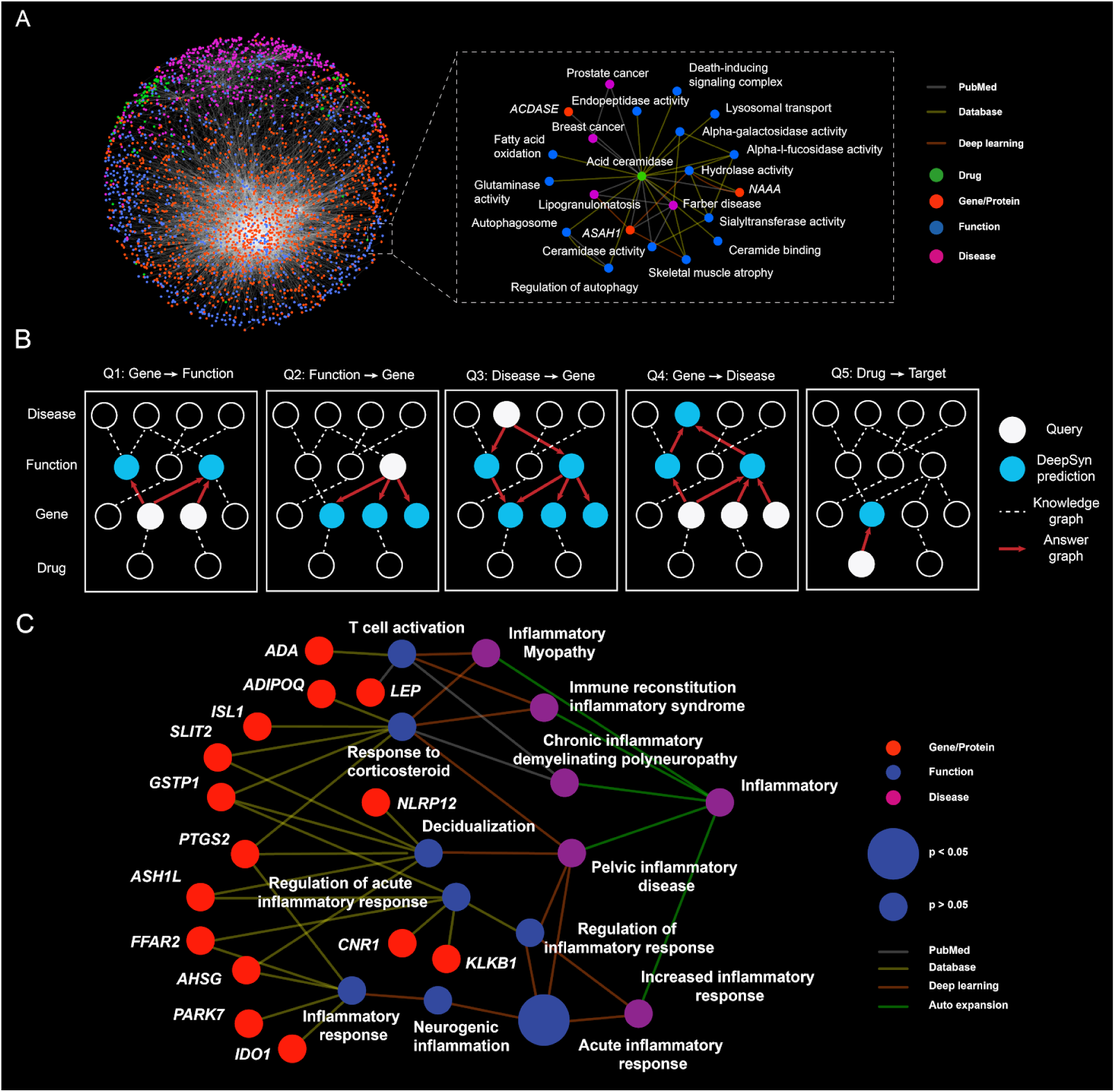
Overview of DeepSyn. **(A)** The DeepSyn knowledge network integrates a wide range of biological databases and biomedical literature. Insets illustrate the direct network neighborhoods of the drug acid ceramidase. **(B)** Five types of prediction. **(C)** The answer graph returned for the query ‘Inflammatory’.

DeepSyn is queried with a set of genes, from which it returns the most relevant functions, diseases, and drugs extracted from the knowledge graph. The query is made context-specific by the addition of conditional constraints, which are also specified in the form of functions, diseases or drugs. Such conditions may even be specified without genes, in which case DeepSyn performs a reverse query to find the most relevant gene set (Figure 1B).

Results are shown in the context of an “answer graph” which is extracted from the global knowledge graph. The answer graph contains direct links from specific to general entities. Each entity in the answer is associated with a *P*-value of significance (the chance that entity would arise from random queries, **STAR Methods**), and each relationship between entities is linked to the supporting evidence in the biomedical literature or databases. For instance, a query with the disease “Inflammatory” produces an answer graph of 8 significantly associated functions and 15 genes (Figure 1C). In constructing the query, DeepSyn automatically adds any additional words and phrases that are frequent in the biomedical literature and contain the query or its conditional constraints. For example, the query “inflammatory” is expanded to add inflammatory myopathy, immune reconstitution inflammatory syndrome, and pelvic inflammatory disease (Figure 1C). Functions associated with more than one inflammatory diseases are in the answer graph, for example the top-ranked gene *PTGS2* regulates the inflammatory response by generating prostaglandins (Hata and Breyer, 2004) and plays a key role in both the treatment of pelvic inflammatory disease (Dhasmana et al., 2014) and immune reconstitution inflammatory syndrome (Shankar et al., 2007).

### Functional synthesis can address multiple types of biological questions

We formulated a panel of gold standard queries to evaluate the accuracy of associating gene sets with functions, diseases, and drugs (Figures 1A-D). The average performance on all of these tasks was relatively high (range 0.74 to 0.96) and represented a substantial improvement in comparison to baseline approaches based on literature co-occurrence and mutual information (**STAR Methods**). First, we examined the ability of DeepSyn to recover the catalog of function names assigned to human gene sets by the Gene Ontology (GO) (Figure 2A). DeepSyn was queried with the set of genes annotated to each GO Biological Process, Molecular Function, or Cellular Component term. For each gene and condition pair, we calculated an empirical *P*-value for the corresponding knowledge graph to represent its significance compared to a random graph (**STAR Methods**). We then looked for terms among all candidate phrases ranked according to the *P*-value to genes in the term. On all three Gene Ontology categories, DeepSyn outperformed both co-occurrence and mutual information-based baseline approaches by at least 18%. Second, we evaluated the reverse function query, in which DeepSyn was given a GO term (function) name and asked to predict the corresponding set of genes. In this case, DeepSyn achieved an average accuracy of 74%, 81% and 78% for BP, CC and MF branches, again significantly outperforming baseline approaches (Figure 2B).

**Figure 2.**
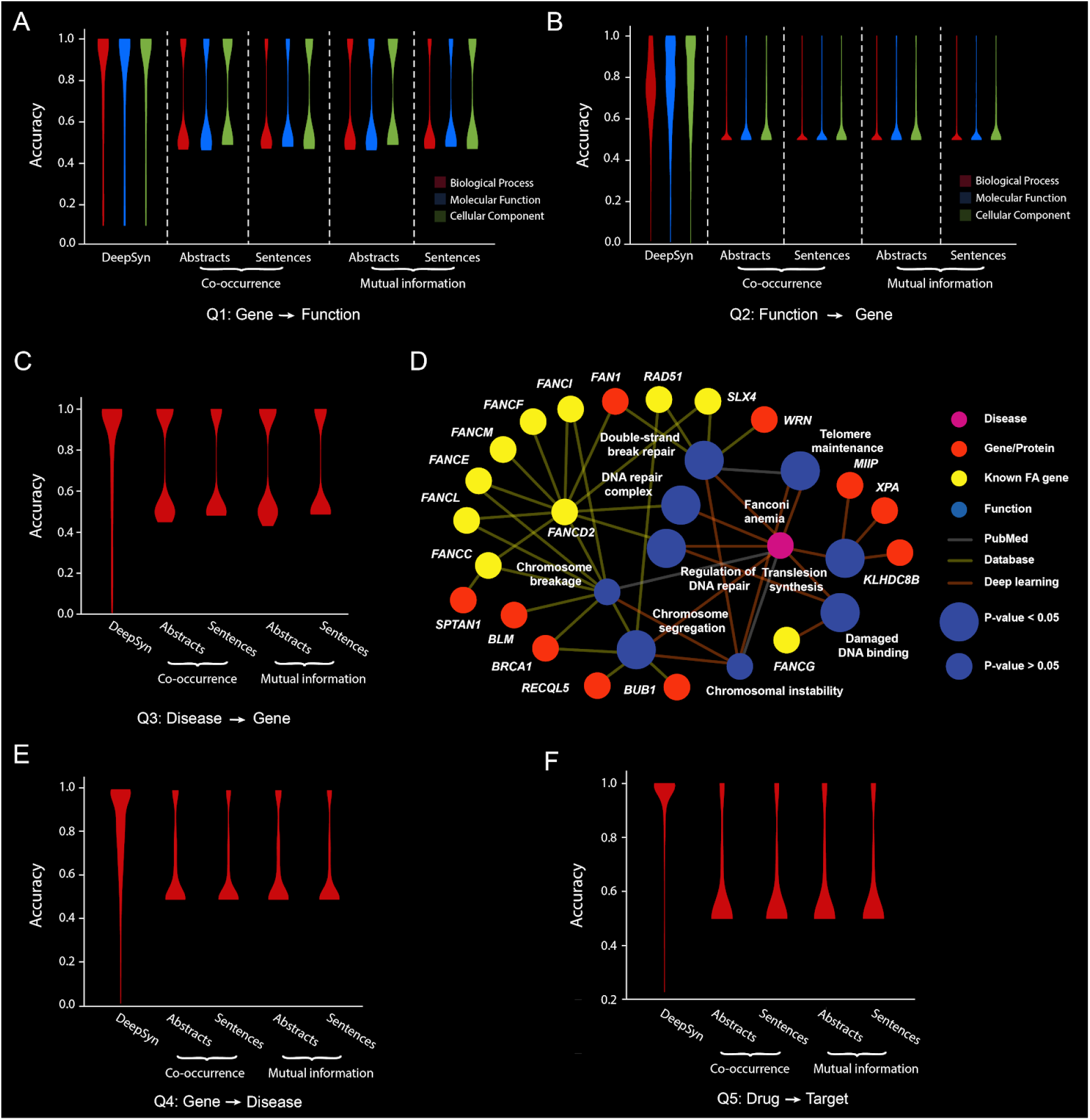
Prediction performance of DeepSyn. **(A)** Accuracy of predicting functions (Gene Ontology terms) for a given gene set. Performance is measured by Area Under Receiver Operating Characteristic (AUROC) curve. **(B)** Accuracy of predicting a gene set for a queried biological function. **(C)** Accuracy of predicting disease genes for a queried disease name. **(D)** Answer graph and associated *P*-value returned for the query ‘Fanconi Anemia’. **(E)** Accuracy of predicting a disease given a gene set. **(F)** Accuracy of predicting targets for a particular drug.

Next, we considered queries related to diseases, i.e. what disease was associated with alterations in a particular set of genes or, alternatively, what genes were associated with a particular disease. Compared to co-occurrence-based methods in the literature, we found that DeepSyn was able to outperform this baseline for at least 21% of the benchmark datasets for these two tasks (Figure 2C). For instance, for the disease query Fanconi Anemia (FA), 11 of the top 20 genes returned in the answer graph had been identified as FA disease genes by previous studies collected by the Monarch database (Köhler et al., 2019) (Figure 2C). The gene *RECQL5* was the top result, which was not documented in Monarch but was found to perform multiple functions in cells defective for the Fanconi anemia pathway (Kim et al., 2015). Further examining the answer graph, we found that *RECQL5* was connected to FA through the function of chromosome segregation (Figure 2D) which is well known to play a role in FA pathogenesis (Cerabona et al., 2014). This link was present in the knowledge graph due to *RECQL5* knockdown experiments, which have shown that *RECQL5* is required for chromosome segregation through interactions with Topoisomerase II α during mid-late S-phase (Ramamoorthy et al., 2012). Thus, DeepSyn was able to identify disease genes not covered by existing biological databases (Haendel et al., 2018). We also evaluated the reverse task, in which DeepSyn was given a set of disease genes and asked to predict the corresponding disease. On a collection of diseases in Monarch, DeepSyn achieved an average accuracy of 0.83 which also showed significant improvement compared to baseline approaches (Figure 2E). Finally, we evaluated the ranking of target genes for a specified drug documented in the DrugBank database (Wishart et al., 2008). We found DeepSyn was able to achieve an accuracy of 96% compared to the baseline approach of 60% (Figure 2F).

### Providing context-specific answers to complex biological questions

As mentioned above, gene set queries can be augmented by the specification of functions, diseases, or drugs under which the gene set has been generated or is otherwise relevant (Figure 3A). Such “conditional” queries can be particularly useful in the functional annotation of gene sets arising from gene or protein expression studies. As a proof of concept, we performed an mRNA expression analysis of 286 glioblastoma tumor samples from The Cancer Genome Atlas (Cancer Genome Atlas Research Network, 2008). The 40 genes with largest variance in gene expression were organized into five clusters based on the cosine similarity (Figure 3B). Notably, none of these clusters (each corresponding to a gene list) was deemed to be significant using standard functional enrichment against the Gene Ontology (**Supplementary Table 1**). In contrast, DeepSyn was able to synthesize significant biological functions for all gene clusters (Figure 3B). For example, the genes *INA, PAN-P2RY11, VSTM2A, ST8SIA3, PCSK2, ACTL6B and CDKN2A* formed a coherent expression cluster across glioblastoma tumor samples. Querying DeepSyn with this gene set and the condition “glioblastoma” produced an answer graph implicating a hierarchy of specific-to-general functions including vessel maturation and autophagy (Figure 3C).

**Figure 3.**
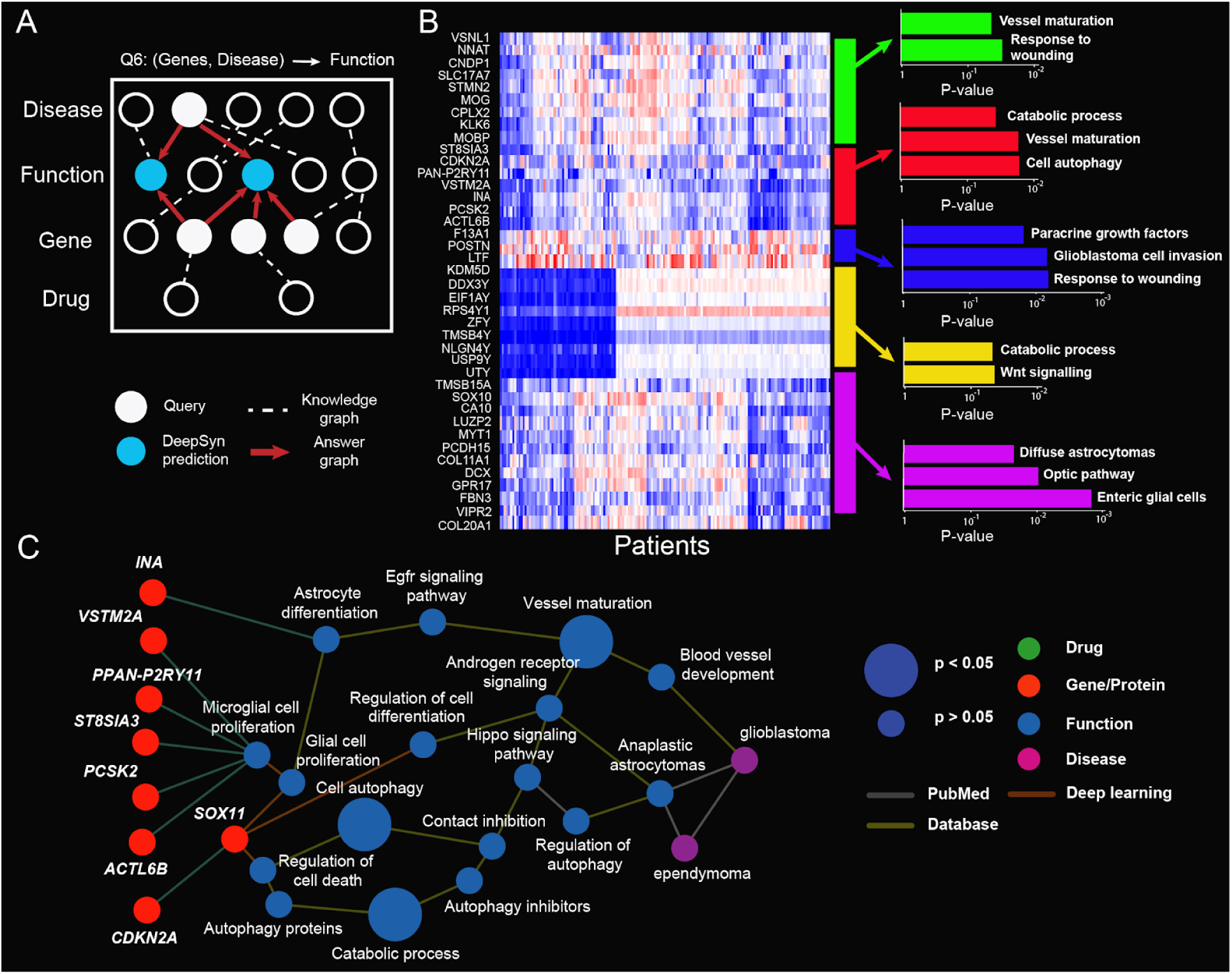
Annotation of genes coordinately expressed in glioblastoma. **(A)** DeepSyn can be used to query a gene set in the context of a disease. **(B)** gene expression clusters identified through analysis of gene expression profiles of glioblastoma tumors (The Cancer Genome Atlas Research Network 2008). Each of the gene clusters defines a gene set and is used to query DeepSyn to identify common functions. The bar plot on the right indicates the *P*-value of each function identified by DeepSyn in the log scale. **(C)** The answer graph returned for one of the glioblastoma gene clusters consisting of seven genes (red clusters in panel **(B)**), with circle size indicating the *P*-value of significance.

Today, gene set enrichment remains the gold standard to measure the success of many biological discoveries in genomics. The main limitation of enrichment methods is that 1) genes perform different functions by interacting with distinct partners in different biological and clinical contexts and 2) they must rely on static manually curated gene functions. To address these limitations, here we have shown that it is possible to use machine learning algorithms to extract context-specific functional information and automatically construct accurate evaluating standards to measure the success rate of biological discoveries. This work enables a philosophical shift in functional analysis, from manual curation of literature to AI-based learning.

## STAR METHODS

### KEY RESOURCE TABLE

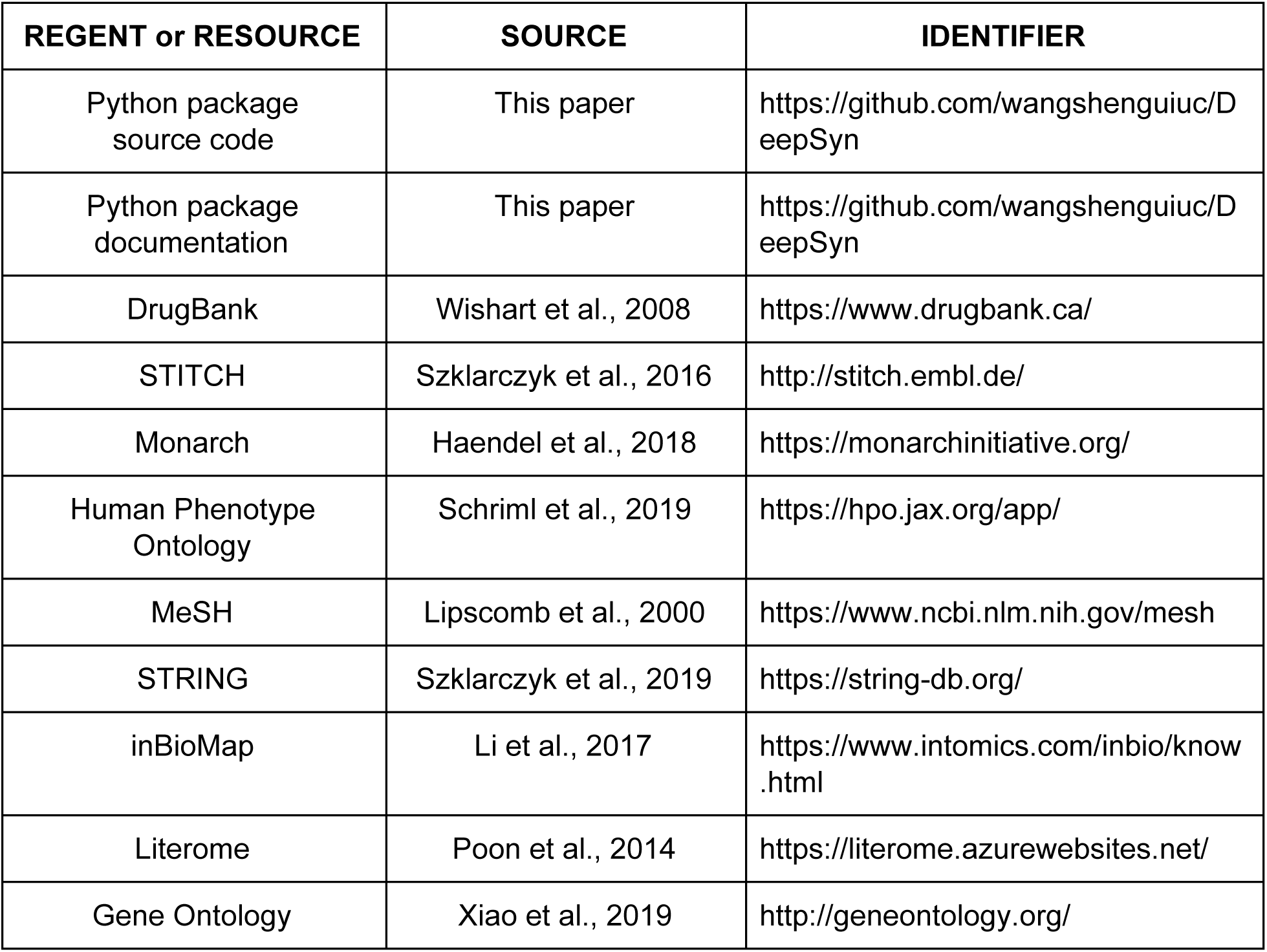

#### Collecting and processing biological databases and literature data

The core of our method is to construct a heterogeneous knowledge graph integrating both literature data and multiple public biological databases. The knowledge graph considers four types of nodes including drug, gene/protein, disease, and biological function. We collected drug-target interactions, disease-disease associations, disease-gene associations, protein-protein interactions, function-gene annotations, and associations between functions. To construct these associations, we collected the following datasets: 1) drug-target interactions from DrugBank (Wishart et al., 2008) and STITCH (Szklarczyk et al., 2016), 2) disease-gene associations from Monarch (Haendel et al., 2018), 3) disease-disease associations from Human Phenotype Ontology (Schriml et al., 2019) and MeSH (Lipscomb, 2000), 4) protein-protein interactions from STRING (Szklarczyk et al., 2019), InBioMap (Li et al., 2017) and Literome (Poon et al., 2014), and 5) function-gene annotations and function-function associations from Gene Ontology (Xiao, 2019). Besides these public databases, associations between two biological entities could also come from PubMed by using text mining techniques. We collected 16,731,155 scientific paper abstracts spanning a wide range of research areas. These abstracts were downloaded using NCBI public APIs (https://www.ncbi.nlm.nih.gov/home/develop/api/). The title of each scientific paper was appended to the corresponding abstract. The median size of each abstract was 200 words and 14 sentences. To mine this large text corpus, we first constructed a vocabulary of high quality phrases (a sequence of one or more words) which was composed from two sources: 1) drug, disease, gene, and biological function names appeared in the database network; 2) phrases mined from PubMed free text by using an unsupervised phrase mining software AutoPhrase (Shang et al., 2018; Wang et al., 2018b). The weight between two phrase nodes was calculated by their mutual information based on their co-occurring probability in the literature. Mutual information is widely adopted in the BioNLP area to handle the frequency bias for calculating the co-occurring probability (Levy and Goldberg, 2014). In addition, we only considered two phrases that co-occurred in >10 articles as literature data was noisier compared to molecular data. The weights of these edges were all normalized to the range of 0 and 1. If there were multiple edges between the same pair of nodes, the maximum value was selected as the final edge weight.

#### Augmenting functional associations by using deep learning

To quantify the relationship between two informative phrases, it is necessary to accurately identify whether a scientific paper is actually relevant to a particular function or not. To address this problem, we trained a neural network model based on the language corpus to explicitly predict whether a scientific paper was associated with a Gene Ontology term or not.

##### Neural network architecture

In this work, we adopted a Convolution Neural Network (CNN) with one convolution layer on top of word embeddings GloVe obtained from an unsupervised neural language model (Pennington et al., 2014). Formally, let *x*_*i*_ ∈ *R*^*k*^ be the *k*-dimensional (*k*=50) word vector corresponding to the *i*-th word in the sentence. The filter output *c*_*i*_ for the *i*-th word was defined as,

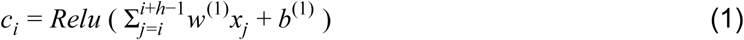

Here *h* was the filter window and was set to 5 in our implementation. *w*^(1)^ was the weight matrix of the linear transformation and *b*^(1)^ was the bias vector. *Relu* was the rectified linear activation function which set values below 0 to 0 (Glorot et al., 2011). We used Eq (1) as a filter to scan the sentence with length *L* and produced a feature map *c* = [*c*_1_, *c*_2_, …, *c*_*L*−*h*+1_]. A max-overtime pooling operation then was applied over the feature map to select the maximum component *c*_max_ as *argmax*{*c*_1_, *c*_2_, …, *c*_*L*−*h*+1_}. The pooling result was then fed into a fully connected layer with a continuous of dropout layer and softmax layer as follows,

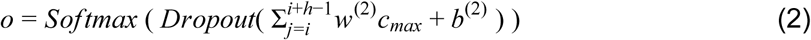

The prediction *o* was associated with a mean-square error loss. *w*^(2)^ and _*b*_(2) were linear weights and bias of this fully connected layer, respectively.

##### Training algorithms and implementation details

Empirically we used the dropout rate of 0.5 and mini-batch size 500. We trained the weights of the neural network by using the gradient descent algorithm ADAM (Kingma and Ba, 2014). The learning rate of ADAM was set as 0.001. The whole computational framework was implemented using the PyTorch library (https://pytorch.org/). In total, we trained 8542 deep neural network models and got an average of AUC value 0.79 evaluated by using 5-fold cross-validation.

##### Training data construction

We constructed a separate learning task for each Gene Ontology term. For each predicting task, we selected all PubMed articles with the term name explicitly showing in abstracts and used these articles as positive training samples. We also selected abstracts that did not contain this term but contained sibling terms (same parent term) on the Gene Ontology as negative training samples. In this way, we encouraged neural network models to learn more discriminative representations of the language corpus by constructing a more challenging machine learning problem. For each term, we randomly select some negative samples to keep its number as ten times of the positive samples.

#### Constructing a biological knowledge graph

This trained CNN classifier was then used to annotate the gene ontology term to each abstract. Such annotation significantly expands existing language models based on co-occurring probability. For instance, an article mentioned “genetic instability” in its abstract and our CNN model also predicted that the article was related to “DNA repair”. In this way, we counted ‘genetic instability’ and ‘DNA repair’ co-occur once. There were two levels of co-occurrence probability: the sentence-level and the abstract-level. The sentence-level co-occurrence probability *P*_*AB*_ between two phrases *A* and *B* was defined as,

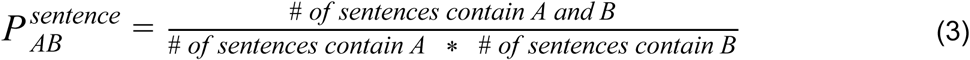

Similarly, the article-level co-occurrence probability was defined as,

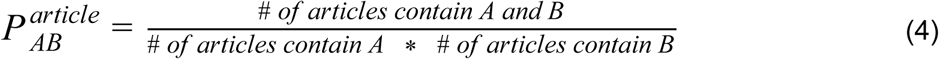

To construct the knowledge graph, we chose to use the article-level co-occurrence probability but we also compared the performance of using sentence-level co-occurrence probability in Figure 2.

#### Querying the biological knowledge graph

Currently, DeepSyn supported four types of user’s queries including drug, disease, gene/protein, and function. Each field could be a set of entities (e.g., a gene set) or empty. If both nodes belong to the Gene Ontology or the Human Phenotype Ontology, we let the more general node point to the more specific node based on the definition of the Gene Ontology and the Human Phenotype Ontology. If two nodes are both informative phrases mined from literature, we let the node with higher occurring frequency in the literature point to the lower frequency one. DeepSyn extracted a subnetwork from the entire knowledge graph with respect to the user’s query by adopting a Depth First Search (DFS) algorithm. The source nodes of the DFS algorithm were the diseases or functions in the user’s query and the target nodes were genes in the user’s query. We also restricted the size of the resulted subnetwork by setting the maximum layer of the network to be less than 5 and each node could be only connected to at most 10 nodes in the network. To calculate the *P*-value for each node in the returned subnetwork, we first summed up the weights of its edges in the subnetwork and then calculated an empirical *P*-value by comparing this score to a background distribution. The background distribution was fit by sampling 10,000 random subnetworks. The *P*-value of a subnetwork was thus defined as the most significant *P*-value of all nodes in this subnetwork. If one of the fields is missing, DeepSyn will automatically search for all the allowed entities in this field and return the union of all the subnetworks based on a *P*-value threshold (=0.05). For instance, if the user only searches a drug’s name ‘gefitinib’ but does not specify any gene names, DeepSyn will automatically generate many queries, such as <“gefitinib”, “*BRAC1*”>, <“gefitinib”, “*TP53*”>…, and union all the returned subnetworks with significant *P*-values.

#### Evaluating the performance of DeepSyn

To evaluate the performance of DeepSyn, we compared DeepSyn with both co-occurrence and mutual information-based baseline approaches. Both approaches were evaluated on sentence and article levels. The sentence-level and article-level mutual information between *A* and *B* were defined as,

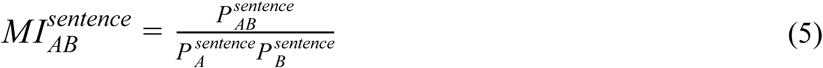

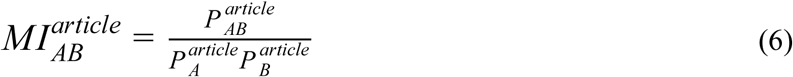

Here 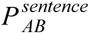 and 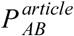 were defined by Eq. (3) and (4). 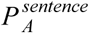 and 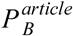 were the probabilities that phrase *A* showed in a sentence or an article, respectively. DeepSyn and all baseline approaches were given a biological query and conditions supported by our system. For example, the user’s query could be a drug name, and all approaches were required to find associated genes. Each approach answered the query by finding the condition that had the highest co-occurrence with the query in the system. DeepSyn and all baseline approaches returned a ranking list of answers. AUROC was then used as the metric to evaluate this ranking list. When our experiments were related to GO, we evaluated the performance on all three categories of GO: Molecular Function, Cellular Component, and Biological Process. We also validated the answer to these queries according to existing biological databases.

## Acknowledgments

This work is supported by NIH LM005652, TR002515, GM102365, the Chan-Zuckerberg Biohub, NIH / NIGMS P41 GM103504, and NIN / NCATS OT3 TR002026.

